# Initiation, elongation and realignment during influenza virus mRNA synthesis

**DOI:** 10.1101/197210

**Authors:** Aartjan J.W. te Velthuis, Judith Oymans

## Abstract

The RNA-dependent RNA polymerase (RdRp) of the influenza A virus replicates and transcribes the viral genome segments in the nucleus of the host cell. To transcribe these viral genome segments, the RdRp ‘snatches’ capped RNA oligonucleotides from nascent host cell mRNAs and aligns these primers to the ultimate or penultimate nucleotide of the segments for the initiation of viral mRNA synthesis. It has been proposed that this initiation process is not processive and that the RdRp uses a prime-realign mechanism during transcription. Here we provide *in vitro* evidence for the existence of this transcriptional prime-realign mechanism, but show that it only functions efficiently for primers that are short or can not stably base pair with the template. In addition, we demonstrate that transcriptional elongation is dependent on the priming loop of the PB1 subunit of the RdRp. We propose that the prime-realign mechanism may be used to rescue abortive transcription initiation events or cope with sequence variation among primers. Overall, these observations advance our mechanistic understanding of how the influenza A virus initiates transcription correctly and efficiently.

**Importance:** The influenza A virus causes severe disease in humans and is considered a major global health threat. The virus replicates and transcribes its genome using an enzyme called the RNA polymerase. To ensure that the genome is amplified faithfully and abundant viral mRNAs are made for viral protein synthesis, the viral RNA polymerase must transcribe the viral genome efficiently. In this report, we characterise a structure inside the polymerase that contributes to the efficiency of viral mRNA synthesis.

## Introduction

RNA viruses use an RNA-dependent RNA polymerase (RdRp) to replicate and transcribe their viral RNA (vRNA) genome. One of the best studied negative strand RNA viruses is the influenza A virus (IAV). The IAV genome is replicated and transcribed in the nucleus of the host cell by the IAV RdRp, an enzyme that consists of the viral proteins PB2, PB1 and PA (1, 2). The N-terminal third of PB2, the PB1 subunit and the C-terminal two-thirds of PA form the conserved core of the RdRp (3-5), while the remaining portions of PB2 and PA form flexible domains that have cap binding and endonuclease activity, respectively (Fig. 1A). Another key functional structure in the RdRp is a conserved PB1 β-hairpin called the priming loop, which resides downstream of the active site of the IAV RdRp and is important for viral replication initiation (4, 6, 7). The promoter for the IAV RdRp is bound by the conserved core of the RdRp and is known to consist of the partially complementary 3ʹ and 5ʹ ends of the viral RNA (vRNA) genome segments (Fig. 1A) (3, 4).

**Figure 1.**
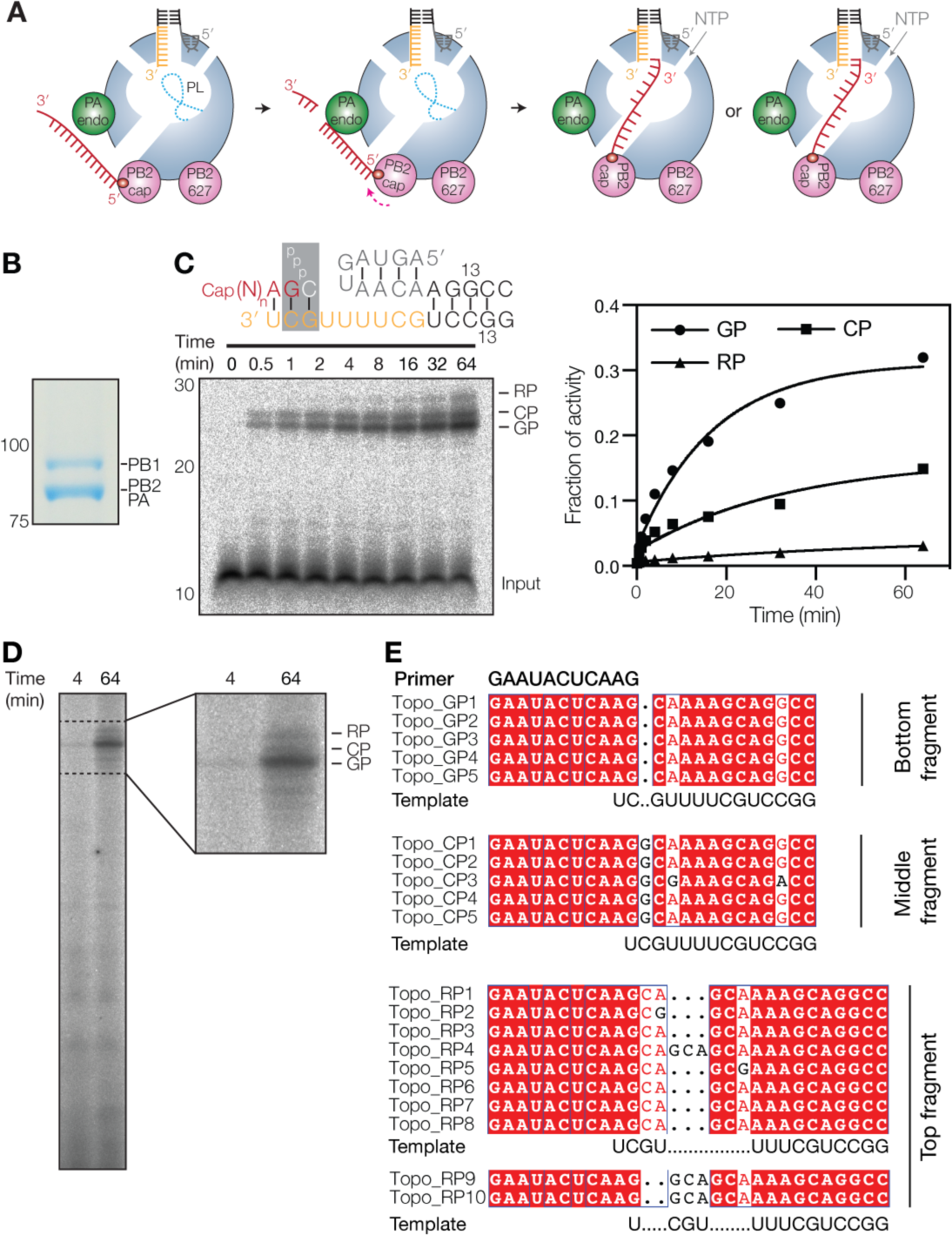
Transcription initiation by the influenza virus RdRp produces three RNA species. (**A**) Schematic of cap cleavage and transcription initiation with a capped 11 nucleotide long primer. The duplex of the viral promoter is shaded black, the 3ʹ end orange and the 5ʹ end grey. The capped primer is shaded red and the priming loop drawn as a dotted line. Alignment of the capped primer to 1U or 2C is also depicted. (**B**) SDS-PAGE analysis followed by Coomassie staining of purified IAV RdRp. (**C**) Extension of a radiolabelled capped 11-nucleotide long primer in the presence of four NTPs. The products produced from position 2C (CP) and 3G (GP) as well as the realignment product (RP) are indicated. The promoter schematic shows the primary initiation site on the wild-type IAV promoter with colours as in Fig. 1A. The graph shows the accumulation of the GP, CP and RP signals as fraction of the total transcriptional activity. Lines represent fits to an exponential decay of one representative experiment. (**D**) Transcription reaction using β-globin mRNA as primer donor. (**E**) Alignment of IAV transcription products present in the GP, CP or RP band containing gel fragments as identified by TOPO cloning and Sanger sequencing.

Unlike IAV replication, in which the RdRp initiates *de novo* (6), the IAV RdRp uses a primer-dependent process for viral transcription initiation. To produce this primer, the IAV RdRp must first bind to the C-terminal domain of an actively transcribing, serine 5-phosphorylated RNA polymerase II (Pol II) complex in the nucleus of an infected cell (8, 9). Subsequent binding and cleavage of nascent Pol II transcripts produces 8-14 nucleotides long capped RNAs (10) (Fig. 1A) that the IAV RdRp can hybridise as primers to the 3ʹ terminus of the vRNA template (Fig. 1A). The PA endonuclease domain has a preference for mRNA cleavage 3ʹ of G residues *in vitro* (11, 12), which creates primers that can be hybridised with the penultimate C (2C) of the 3ʹ terminus of the vRNA template (Fig. 1A). This match between cleavage preference and template sequence is also reflected in a recent crystal structure of the influenza B virus RdRp that is bound to a vRNA and capped primer (13), because it showed that the 3ʹ terminus of the vRNA can ‘overshoot’ the active site by 1 nt without duplex unwinding. Hence, ostensibly by the default, the RdRp positions 2C of the vRNA in the -1 position of the active site (13), which is ideal for transcription initiation with primers ending in 3ʹ G from 3G of the template. However, in viral infections and other *in vitro* studies, capped RNA primers with other 3ʹ terminal bases are also frequently produced and used (14-18) and current evidence suggests that these primers are extended from 2C instead of 3G (Fig. 1A).

After transcription initiation, the RdRp extends the primer in a template-dependent fashion. However, IAV mRNAs isolated from infected cells often contain 3-nt repeats (GCA or GCG, depending on the segment) that are complementary to the 2^nd^, 3^rd^ and 4^th^ nucleotide of the template (14, 15). This observation implies that RdRp processivity is limited over the first 4 bases of the vRNA. It has been proposed that the 3-nt repeats are introduced by the IAV RdRp through a realignment mechanism (14, 15, 18), but direct evidence for this process is currently lacking. Moreover, it is also not known whether there is a link between the generation of capped primers in the host nucleus, the ability of the RdRp to hybridise these primers efficiently to the 3ʹ terminus of the vRNA template, and the generation of these 3-nt duplications.

In this study, we use a combination of structure-guided mutagenesis and *in vitro* polymerase activity assays to provide *in vitro* evidence for the existence of low-processive transcription elongation events that result in a duplication of the first three nucleotides of the vRNA 3ʹ terminus. Moreover, we show that the synthesis of these duplications is dependent on the sequence of the capped primer, the sequence of the template, and the interaction of the body of the RNA primer with the body of the priming loop. These observations thus provide a mechanistic insight into IAV RNA synthesis and redefine the function of the priming loop as a platform for both efficient replication (6, 7) and transcription.

## Results

### The initiation of IAV transcription produces multiple products

IAV transcription uses a capped RNA primer that is snatched from host cell mRNAs and subsequently hybridised to the 3ʹ 1U and/or 2C of the vRNA promoter for extension from 3ʹ 2C or 3G, respectively (Fig. 1A). Although it is currently assumed that this process is largely dependent on Watson-Crick base pairing between the primer and the template, transcription initiation without Watson-Crick base pairing has been observed (14-18). To study IAV transcription initiation in detail, we expressed the PB1, PB2 and PA subunits of the influenza A/Northern Territories/60/1968 (H3N2) virus in insect cells (9) and purified the recombinant enzyme using a tandem affinity purification (TAP) tag on PB2 as described previously (5). The purified enzyme was analysed by SDS-PAGE for purity (Fig. 1B). Next, we setup reactions containing the IAV RdRp with an 11-nt radiolabelled capped RNA ending in 3ʹ AG (AG primer) and followed the extension of the primer in time (Fig. 1C). PAGE analysis showed that extension of the primer resulted in one major product and two slower migrating minor products (Fig. 1C). The slowest migrating product was produced ~10 times less efficiently than the major product (Fig. 1C). Transcription products similar to the ones described above have previously been observed in assays in which IAV RdRp purified from mammalian cells was used to extend a 11-nt long capped primer (6, 7) or a β-globin mRNA-derived primer (19-23) (Fig. 1D), suggesting that these three RNA species are typical IAV transcription products.

To characterise the three products in more detail, we cut the part of the gel containing the three bands into three and extracted the RNA contained in the three gel fragments. Subsequent Sanger sequencing analysis revealed that the gel fragment with the bottom band contained an RNA species that had initiated at 3ʹ 3G of the template (Fig. 1E), whereas the fragment with the second band contained a product that had been produced after initiation at 3ʹ 2C of the template (Fig. 1E). The third fragment, which included the third band and a small part of the gel above this band, was found to contain three RNA species (Fig. 1E) that all had at least one duplication of residues 2-4 of the template, similar to the RNA species observed in IAV infections (14, 15, 18). Moreover, the RNA species contained products that had initiated from both 3ʹ 2C and 3ʹ 3G, implying that the duplication of residues 2-4 during transcription elongation occurs independently of the site of transcription initiation. From here on, we will refer to the identified RNA species as the 3G initiated product (GP), the 2C initiated product (CP) and the 3-nucleotide repeat products (RP) for simplicity.

### Realignment during transcription is dependent on 3ʹ 1U and 4U of the vRNA template

To verify that the GP species had indeed been synthesised after initiation at 3ʹ 3G in our assay, we first mutated the 3ʹ 2C of the vRNA promoter to A (2C→A) in order to disrupt G-C base pairing between the AG primer and template. As shown in Fig. 2A, this mutation resulted in a loss of the GP signal and a concomitant increase in the CP signal (Fig. 2A), likely due to an increase in G-U base pairing between the AG primer and template (note that the CP band generated on the mutant template migrates between the GP and CP bands that were produced on the wild-type template. This difference is due to a G to U change, which reduces the molecular weight of the CP band and increases its mobility in 20% PAGE). In the second control, we replaced the AG primer with a primer ending in 3ʹ CA and incubated this with the wild-type promoter, NTPs and the RdRp. As shown in Fig. 2B, this reaction yielded a similar CP product level, confirming that GP synthesis is dependent on primer base pairing with 3ʹ 2C and initiation at 3ʹ 3G. A similar result was obtained when we used a primer ending in 3ʹ AA (Fig. 2C). To fully verify that CP synthesis is dependent on base pairing with 1U, we next mutated 3ʹ 1U of the vRNA promoter to A (1U→A) and incubated this promoter with the AG primer and the IAV RdRp. We found that in this reaction the CP as well as the RP level had both been significantly reduced (Fig. 2D and E), which confirms that CP synthesis is dependent on primer base pairing with 3ʹ 1U. Interestingly, when we incubated the CA or the AA primer with the 1U→A template, we found that transcription initiation had been severely impaired, suggesting that A-A base-pairs do not support efficient transcription initiation.

**Figure 2.**
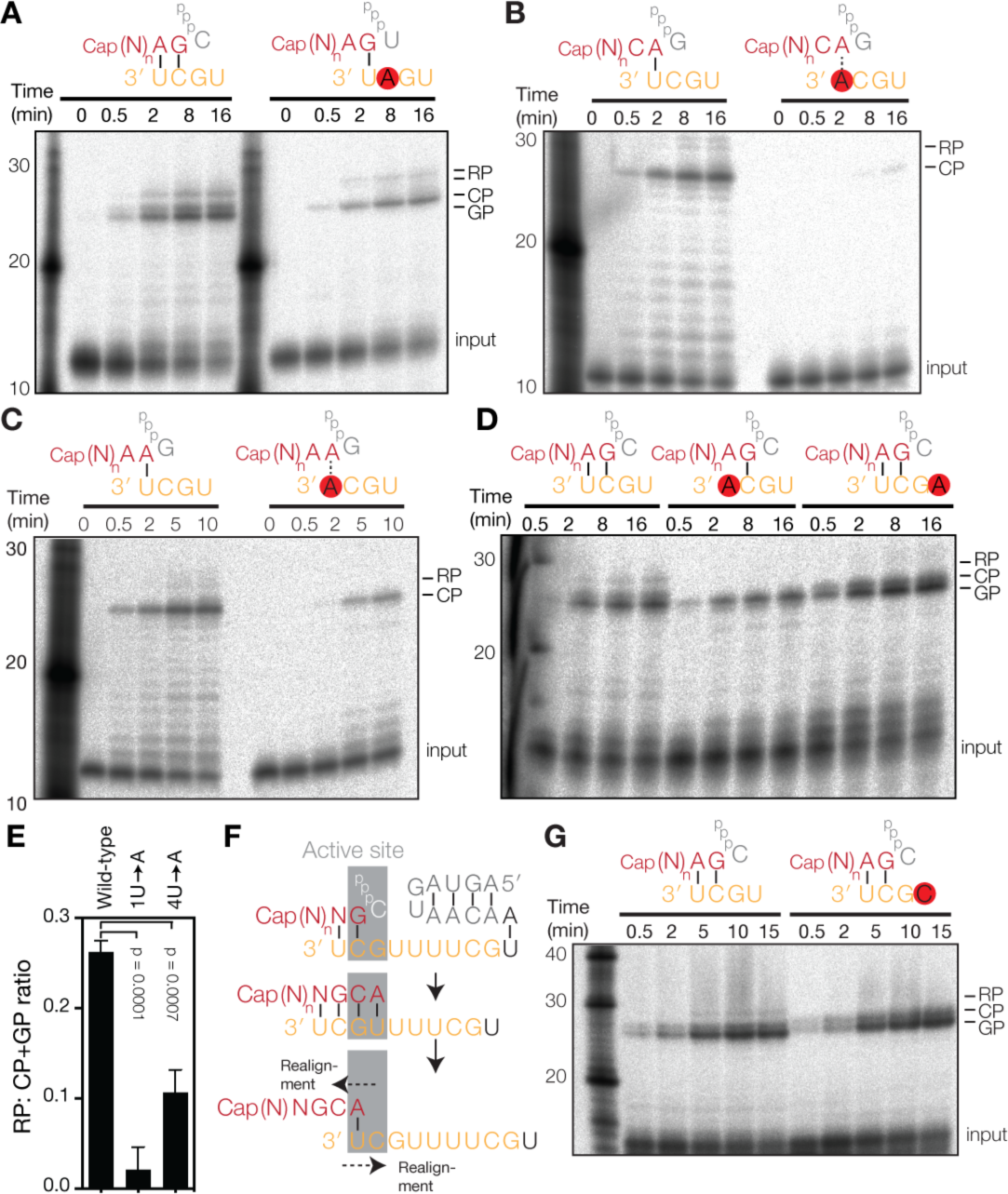
Realignment during IAV transcription involves 4U and 1U of the template strand. (**A**) Extension of a radiolabelled capped 11-nucleotide long RNA in the presence of unlabelled NTPs and IAV RdRp. The schematic shows the primary initiation site on the wild-type or mutant 2C→A 3ʹ promoter strand. (**B**) Extension of a radiolabelled capped 11-nucleotide long RNA primer ending in 3ʹ CA on the wild-type vRNA promoter or the 3ʹ 1U→A mutant promoter. (**C**) Extension of a radiolabelled capped 11-nucleotide long RNA primer ending in 3ʹ AA on the wild-type vRNA promoter or the 3ʹ 1U→A mutant promoter. (**D**) Extension of a radiolabelled capped 11-nucleotide long RNA primer ending in 3ʹ AG on the wild-type, 3ʹ 1U→A, or 3ʹ 4U→A vRNA promoter. (**E**) Quantitation of Fig. 2D. Graph shows mean RP: GP+CP ratio of 3 independent assays. Error bars indicate standard deviation. The p-values were determined using an unpaired t-test. (**F**) Model of realignment after initiation from 3G. Colours as in Fig. 1A. (**G**) Extension of a radiolabelled capped 11-nucleotide long RNA primer ending in 3ʹ AG on the wild-type vRNA promoter or a 3ʹ 4U→C mutant promoter.

In the above experiments, RP formation was independent of 3ʹ 2C, but dependent on 3ʹ 1U (Fig. 2D). This is in line with the idea that the RP species are formed through a realignment event between positions 4U and 1U (Fig. 2F). To confirm this, we mutated 4U to A (4U→A) in the 3ʹ strand of the vRNA promoter and found that this mutation supported CP and GP formation, but greatly reduced RP synthesis (Fig. 2D). Indeed, quantitation of the product levels showed that the ratio between the RP signal and the sum of the CP and GP (CP+GP) signals was significantly diminished (Fig. 2E). A similar result was obtained when we mutated 4U to C (4U→C) (Fig. 2G), a sequence variant that down-regulates transcription of segments 1, 2 and 3 of the IAV genome (24). Overall, these findings imply that Watson-Crick base pairing between the primer and the template plays an important role during transcription initiation and that Watson-Crick base pairing between the extended primer and 3ʹ 1U is crucial for RP formation when transcription elongation is not processive.

### Duplex stability and ssRNA primer length affect transcription processivity

The above results imply that transcription initiation relies on base pairing when the IAV RdRp uses primers ending in AG or CA. However, the IAV RdRp can also use primers that are not complementary to the template (15, 16, 18). To investigate how such non-complementary primers affect transcription initiation and elongation processivity, we first replaced the terminal G of the AG primer with U (i.e. creating a primer ending in 3ʹ AU; AU primer). On the wild-type vRNA promoter, this AU primer was efficiently extended into a major CP signal and minor GP and RP bands (Fig. 3A). To confirm that the AU primer was indeed primarily extended from 3ʹ 2C, we replaced the wild-type promoter with the 1U→A mutant promoter, because this would enforce U-A base pairing with the 3ʹ A of the mutant promoter and increase the CP:GP signal ratio. Indeed, the control reaction yielded a similar CP signal and a weak GP band (Fig. 3A). The RP signal has been severely reduced in this reaction, due to the 3ʹ 1U to A mutation in the template, which prevented realignment during non-processive elongation. Thus, we find that U-U base pairs support transcription initiation, while A-A base pairs do not (Fig. 2B and C).

**Figure 3.**
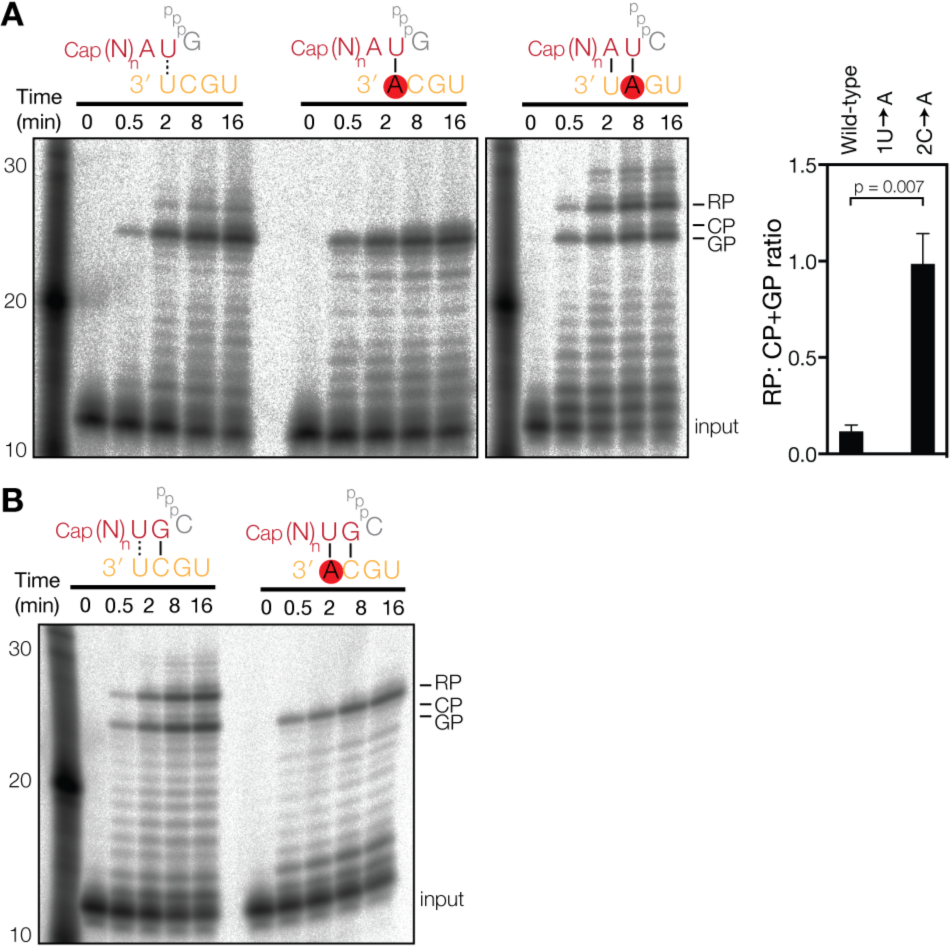
Transcription elongation is affected by the primer sequence. (**A**) Extension of a radiolabelled capped 11-nucleotide long RNA with 3ʹ AU sequence in the presence of four NTPs. Graph shows mean RP: GP+CP ratio of 3 independent assays. Error bars indicate standard deviation. The p-values were determined using an unpaired t-test. (**B**) Extension of a radiolabelled capped 11-nucleotide long primer with 3ʹ UG sequence on the wild-type vRNA promoter or the 3ʹ 1U→A mutant promoter.

To investigate if we could make the AU primer initiate from G3, we next used the 2C→A mutant promoter in the reaction (Fig. 3A) and found that A-U base pairing at positions 1 and 2 of the primer-template duplex supports GP formation. Surprisingly, this altered initiation duplex also upregulated realignment (Fig. 3A) relative to the wild-type template and the 3ʹ AG primer reaction (Fig. 1C). In addition, we observed multiple RP bands of higher molecular weight (Fig. 3A), in line with the multiple RP sequences identified in Fig. 1E and elsewhere (15). Together, this suggests that the duplex that is formed between the 2C→A mutant promoter and the AU primer is less stable and that this reduces the processivity of the IAV RdRp. However, the difference in stability among the various primers and promoters tested is minimal, in particular when we take into account their partial extension products. For instance, the duplex formed during extension of the AU primer on the wild-type promoter is 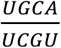, whereas the duplex is 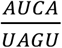 when the AU primer is extended on the 2C→A mutant promoter. We therefore reasoned that the stability of the duplex could not solely explain the differences in processivity and we suspected that the ssRNA length of the primer (i.e., the number of bases that is not bound by the template) could play a role in RP synthesis as well.

To investigate this further, we performed a transcription reaction with a primer ending in 3ʹ UG (UG primer). Up to the point of realignment, this primer is extended into the same 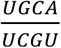 primer-template duplex as the 3ʹ AU primer on the wild-type template. However, because the UG primer only has 9 ssRNA bases after annealing to the 3ʹ 2C of the promoter (compared to 10 ssRNA bases for the AU primer after annealing with 3ʹ 1U), we would expect an effect on RP formation. Indeed, during extension of the UG primer equal levels of GP and RP were formed (Fig. 3B). Moreover, RP formation was abolished by mutation of 1U (Fig. 3B) without a change in the GP signal level, confirming the identity of the RP band. This thus demonstrates that IAV transcription processivity is affected by both the stability of the primer-template duplex and the ssRNA length of the primer.

### The priming loop suppresses realignment during transcription

The above observation suggests that a component of the RdRp interacts with the primer to suppress realignment and increase processivity. Superposing the influenza B virus RdRp with bound primer onto the poliovirus 3D^pol^ elongation complex shows that the PB1 priming loop, which is located between the active site and the entrance of the nascent strand exit channel, is ideally positioned to interact with the incoming capped primer (Fig. 4A). The PB1 priming loop was previously found to be crucial for viral replication initiation and prime-realignment (6, 7). Moreover, it was shown that deletion of residues 648-651 of the priming loop (Δ648-651) increases IAV transcription (6), which suggests that the priming loop may play a role in transcription initiation or elongation. To test whether the priming loop affects IAV transcription elongation, we purified a set of seven priming loop truncation mutants alongside an active site and wild-type control (Fig. 4B and C) as described elsewhere (7), and incubated these with the AG primer and NTPs. We observed clear GP and CP signals in six of the nine reactions as well as a double RP band, likely because both GP and CP initiation products had been realigned (see also Fig. 3A). Measurement of the synthesised CP and GP signals showed that mutant Δ648-651 and a mutant lacking PB1 residues 642-656 (Δ642-656) extended the capped primer more efficiently than the wild-type enzyme (Fig. 4C and D). This suggests that in the wild-type enzyme, the tip and β-sheet of the priming loop affect elongation of the capped RNA primer. By contrast, mutant Δ636-642 synthesised CP and GP levels that were indistinguishable from wild-type, while the four other priming loop mutants showed impaired primer extension levels (Fig. 4C and D), likely because they had a general activity impairment (7).

**Figure 4.**
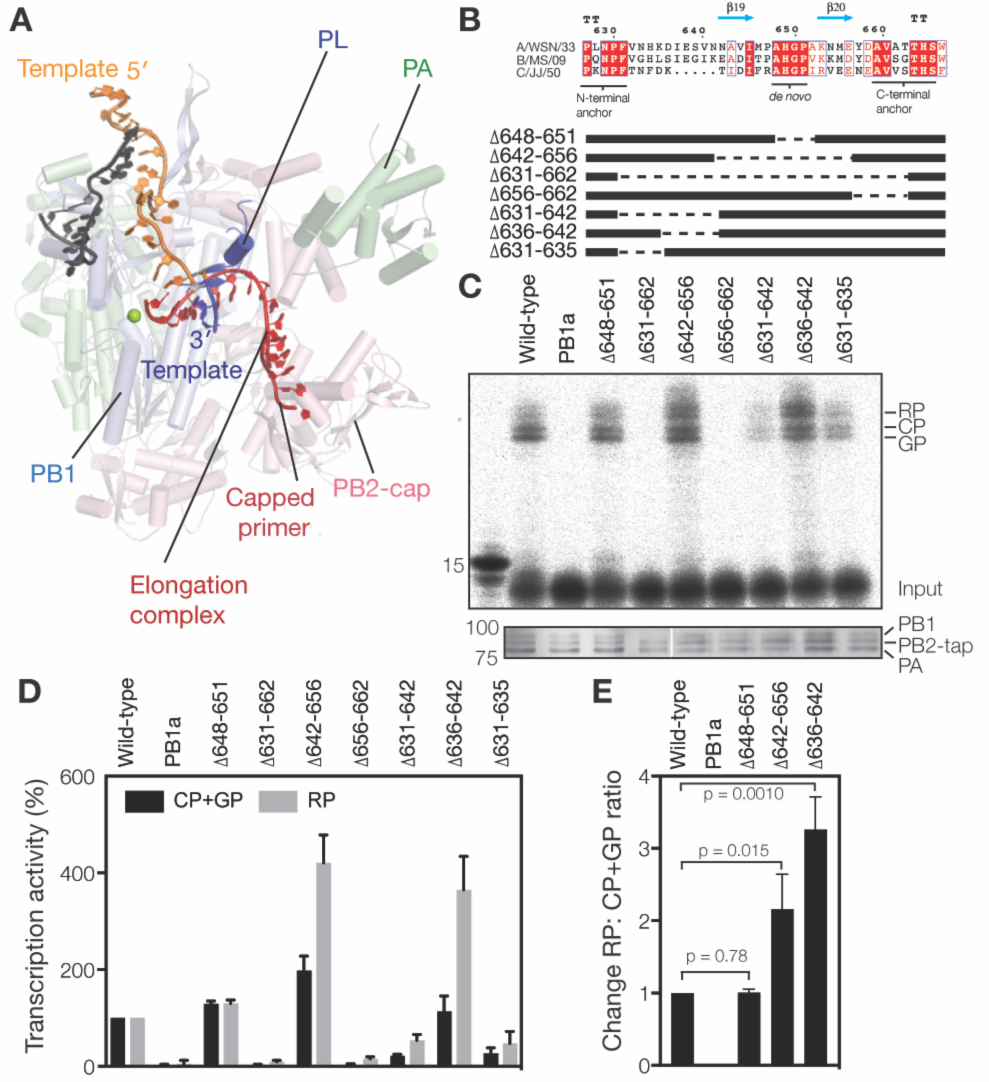
The priming loop affects transcription elongation. (**A**) Superposed structures of the influenza B virus RdRp (PDB 5MSG) with the poliovirus 3D^pol^ RdRp elongation complex (PDB 3OL7). Of the poliovirus 3D^pol^ complex, part of the nascent strand (red) and two nucleotides of the template (dark blue) are shown. These superposed structures illustrate the putative path of the partially resolved capped primer (red) from the 5MSG structure and the position of the priming loop (PL) relative to the putative path of the rest of the capped primer. (**B**) Amino acid alignment of the PB1 priming loop sequences of the IAV A/WSN/33 (H1N1), influenza B virus B/Michigan/22687/09, and influenza C virus C/JJ/50. Identical residues are shaded red, conserved residues boxed in blue and the secondary structure interpretations based on PDB 4WSB. Also shown is a schematic of the PB1 priming loop truncation mutants (7) used in this study. (**C**) Extension of a radiolabelled capped 11-nucleotide primer ending in 3ʹ AG in the presence of wild-type or mutant IAV RdRp purified from 293T cells. Bottom panel shows SDS-PAGE analysis followed by silverstaining of purified IAV RdRps. (**D**) Mean accumulation of transcription products initiated from 3G and 2C (GP+CP) or produced after realignment (RP). Error bars show standard deviations (*n* = 3). (**E**) Fold change in realignment relative to wild-type after normalisation to the total transcription activity. Error bars show standard deviations (*n*= 3). The p-values were determined using an unpaired t-test.

We next analysed the effect of the priming loop on transcription elongation in more detail and measured the RP signal produced by the mutants. This analysis showed that transcriptional realignment appeared increased in reactions containing mutants Δ642-656 and Δ636-642 (Fig. 4C-E). After correction for the differences in the transcriptional activity, we found that the realignment efficiency of mutants Δ642-656 and Δ636-642, as expressed as the change in RP: CP+GP ratio, was significantly increased compared to wild-type, whereas Δ648-651 was indistinguishable from wild-type (Fig. 4E). These results imply that the β-strand of the priming loop suppresses realignment during transcription elongation and thus plays a role in the processivity of IAV transcription. Due to the absence of sequence conservation in the β-strand of the loop (Fig. 4B), future crystallographic studies will be required to identify the interaction mechanism between the primer and the priming loop.

## Discussion

The mechanism of influenza virus transcription relies on binding and cleavage of capped host cell mRNAs and the alignment of the cleavage products to the vRNA template (Fig. 1A). Cleavage is mediated by the PA endonuclease, which preferentially cleaves 3ʹ of G moieties *in vitro* (11, 12). A crystal structure of the influenza B virus RdRp bound to a vRNA template and a capped primer showed that the 3ʹ terminus of the vRNA can enter and even ‘overshoot’ the active site by 1 nt without duplex unwinding (13). This places 2C and 3G in positions -1 and +1 of the active site (Fig. 5), ideal for transcription initiation with primers ending in 3ʹ G. We here find that primers ending in 3ʹ G are indeed preferentially base paired with 2C of the template to allow initiation from 3G. However, a substantial fraction of the primers ending in G is also base paired with 1U, enabling initiation from 2C and suggesting that although Watson-Crick base pairing is important for initiation, Watson-Crick base pairing is not essential. Indeed, we also observe that U-U base pairs support efficient initiation, although A-A base pairs do not, which suggests that transcription initiation is subject to additional constraints to base pairing, such as the shape of the primer-template helix.

**Figure 5.**
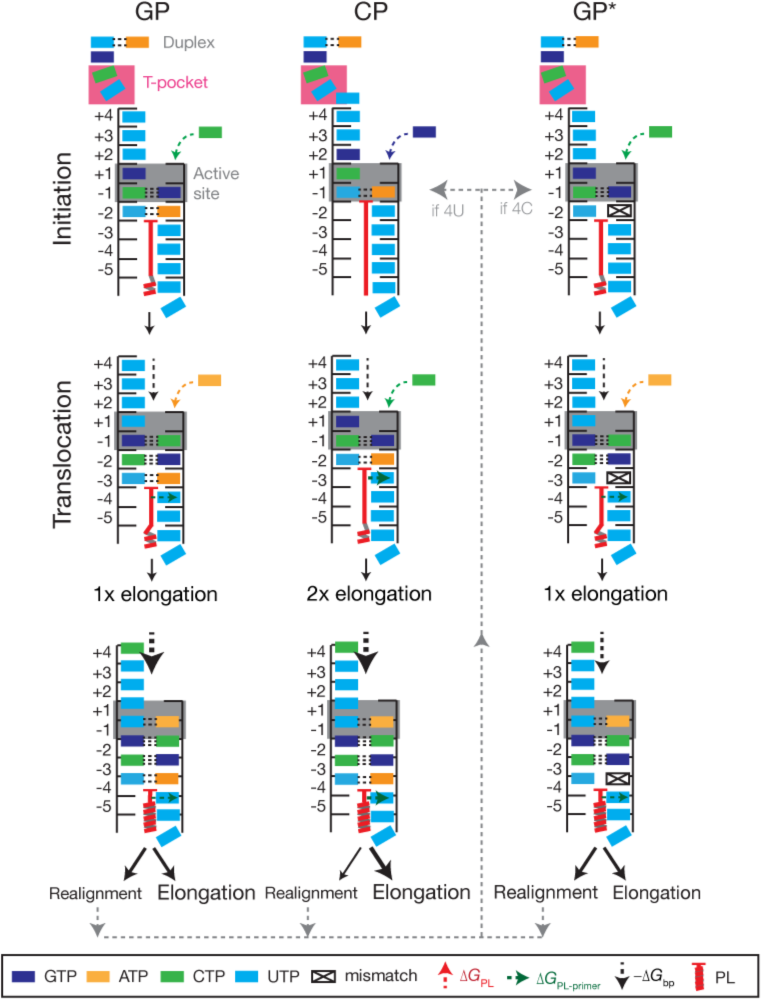
Model for influenza A virus transcription initiation and elongation. IAV transcription can initiate with a capped primer that fully base pairs with the 3ʹ 1U and 2C of the vRNA (GP formation), a primer that base pairs only with the 3ʹ 1U (CP formation) or a primer that only Watson-Crick base pairs with 2C and not 3ʹ 1U (GP* formation; non-Watson-Crick base pairing is indicated as a mismatch). Based on PDB 5MSG, 6 bases of the 3ʹ vRNA terminus are located in the template entry channel during GP synthesis, while three bases (3ʹ 7U, 8C and 9G) remain single-stranded between the duplex at the surface of the RdRp and the template entry channel. Of these three residues, 7U and 8C are stacked in a T-orientation by residues of the PB2 subunit (the ‘T-pocket’). It is likely that the interaction between PB2 and 7U is sequence specific as a 7U→A mutation was previously shown to abrogate *de novo* initiation (29). The priming loop stabilises the capped primer in the active site (Δ*G*_PL-_ primer) to suppress realignment. When the 3ʹ 1U of the vRNA leaves position -4 of the active site, realignment may occur depending on the stability of the template-primer duplex. On a vRNA promoter containing a 3ʹ 4U, realignment will proceed (grey dotted line) via the CP pathway. On a vRNA promoter containing a 3ʹ 4C (segment 1, 2, and 3 of the IAV genome), realignment will proceed via the GP* pathway. It is known that this mutation down regulates transcription (24).

When no G is available in the first 14 nt of the capped host mRNA, PA-mediated cleavage can occur at other bases *in vitro* (11, 12) and *in vivo* (15, 17). We find that primers ending in a 3ʹ A or U are preferentially aligned to 1U of the template and extended from 2C (Fig. 2 and 3). We did not analyse transcription initiation from primers ending in 3ʹ C in this study. To support transcription initiation from 2C, residues 1U and 2C of the vRNA 3ʹ terminus must be placed at positions -1 and +1 of the active site, respectively (Fig. 5). If initiation from 3G is the default for the RdRp (13), initiation from 2C can occur only if the 3ʹ terminus backtracks 1 nt. This step would leave 5 nt in template entry channel and 1 unpaired base upstream of the entry channel (Fig 5). Although it is presently unknown where this unpaired base can be harboured in the structure, it is tempting to speculate that it can be accommodated below bases 7U and 8C, which are stacked in a T-orientation by residues of the PB2 subunit (the ‘T-pocket’). Given that we observe significant initiation from 2C with primers ending in G (Fig. 1C), backtracking of the template is likely a frequent event. An alternative explanation is that initiation from 2C is the default initiation position and that the template must track forward to facilitate initiation from 3G, but this notion is not supported by crystal structures at present.

Previous analysis of IAV mRNAs from infected cells identified GCA duplications in the 5ʹ termini of these mRNAs (15, 18). Our *in vitro* analysis of IAV transcription elongation suggests that these 3-nt duplications are the result of a realignment event that shifts a partially extended capped primer from residue 4U of the vRNA 3ʹ end to 1U (Fig. 2F, 5). It is unlikely that the duplications are generated through cleavage of (partly) finished IAV transcription products and their subsequent re-extension into full-length transcription products. First, we observe that the formation of the GCA duplications is dependent on template residues 1U and 4U (Fig. 2E and F) and not on 2C (Fig. 3A). The latter base allows the incorporation/presence of a G in the transcript and this would be the preferred site for PA endonuclease cleavage (11, 12) over cleavage downstream of the A in the GCA sequence. Second, there is no clear evidence that cleavage products accumulate in reactions that rely on internal radioactive labelling, such as the globin-primed transcription reactions (Fig. 1D). Lastly, an extension-cleavage-extension reaction is likely more complicated than the proposed realignment mechanism, because it would require removal of the partially extended primer from the product exit channel, a conformational change of the PB2 cap binding domain to allow primer egress, cleavage of the transcript by the PA endonuclease, and reinsertion of the primer. Overall, we thus conclude that a realignment mechanism is the most parsimonious explanation of the findings reported here and in the literature.

In the wild-type RdRp, realignment events appear to be relatively rare (~10% of elongation events) when the RdRp is extending a primer that can form a double Watson-Crick base pair with the template (i.e., the AG primer). However, 15 other nucleotide combinations exist for the last 2 bases of the primer, which implies that realignment may be more frequent in viral infections. Indeed, in viral infections up to 30% of transcription elongation events involve realignment as shown by deep-sequencing (15). Although we do not understand the importance of the realignment mechanism in IAV RNA synthesis, we do observe an increase in realignment events when i) primer and template are not aligned through Watson-Crick base pairing and ii) the part of the primer that does not base-pair with the template is relatively short (Fig. 3). Without the realignment mechanism, transcription initiation in the above 2 scenarios would produce significantly fewer full-length IAV mRNAs (on a UG primer ~50% of elongation products is realigned; Fig. 3B), which would limit viral protein synthesis in turn. We therefore speculate that the realignment mechanism is a means to rescue low-processive transcription events and limit abortive transcription initiation. However, the virus may also use this mechanism to regulate the expression of its genes. Indeed, segments with a 3ʹ 4C in the vRNA promoter instead of the canonical 3ʹ 4U show little realignment (Fig. 2G), which may explain why transcription is down regulated on these IAV genome segments (24).

In summary, we here propose that the priming loop of the IAV RdRp interacts with the primer that it snatched from the host cell to improve transcription elongation, which, as we argued above, may be crucial for IAV mRNA and protein synthesis. This thus provides the IAV RdRp with two mechanisms to optimise viral transcription and help the enzyme elongate efficiently after a cap-snatching event. These observations provide a deeper insight into IAV RNA synthesis and they may be relevant for other negative strand RNA viruses that have been observed to use realignment mechanisms as well.

## Experimental procedures

### Cells and plasmids

Human embryonic kidney (HEK) 293T cells were maintained in DMEM (Sigma) supplemented with 10% fetal calf serum (FCS). Sf9 cells were grown in XPRESS medium (Lonza) supplemented with penicillin and streptomycin. Plasmids pPolI-NA, pcDNA-NP, pcDNA-PB1, pcDNA-PA, pcDNA-PB2-TAP, and pcDNA-PB1a have been described previously (19, 20, 25). Also the priming loop mutant PB1 Δ648-651 (6) and the PA endonuclease mutant D108A (26) and priming loop mutants Δ642-656, Δ636-642, Δ631-662, Δ656-662, Δ631-642 and Δ631-635 have been described elsewhere (7).

### Sequence alignment and structural modelling

Amino acid sequences of the PB1 subunits of IAV A/WSN/33 (H1N1), influenza B virus B/Michigan/22687/09, and influenza C virus C/JJ/50 were aligned using ClustalX (27). The alignment was visualised using ESPript (28). To superpose the poliovirus 3D^pol^ elongation complex (PDB 3OL7) and the influenza B virus RdRp crystal structure (PDB 5MSG), we aligned active site residues 324-332 of the poliovirus enzyme with residues 442-449 of the bat influenza virus PB1 subunit in Pymol 1.3.

### Capping and labelling RNA primers

Synthetic 5’ tri- or diphosphate-containing RNAs of 11 nt (Table 1, Chemgenes) were capped with a radiolabelled cap-1 structure using 0.25 µM [α-^32^P]GTP (3,000 Ci mmole^−1^, Perkin-Elmer), 2.5 U/µl 2’-O-methyltransferase (NEB) and a vaccinia virus capping kit (NEB). The products were denatured in formamide and purified as described previously (6).

**Table 1:**
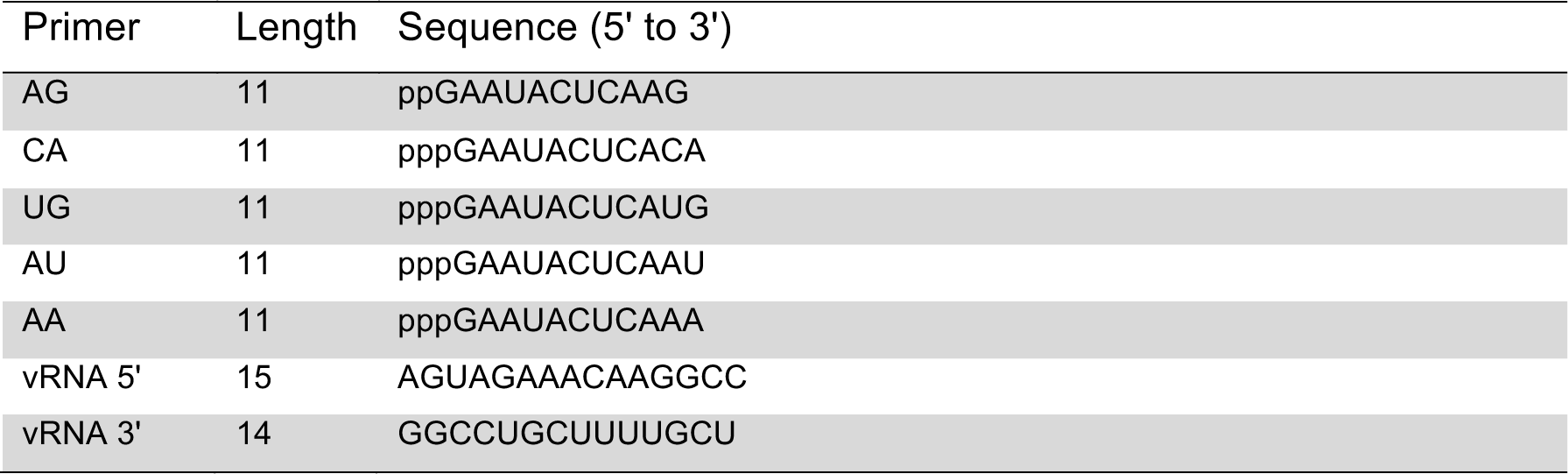
RNA primers.

### *In vitro* capped oligo nucleotide extension assay

Capped RNA oligo extensions were typically set up as 25-μl reactions containing: 1 mM DTT, 5 mM MgCl_2_, 1U/μl RNAsin (Promega), 1500 cpm capped RNA primer, 0.7 μM vRNA promotor (Sigma), 5% glycerol, 0.05% NP-40, 75 mM NaCl, 10 mM HEPES pH 7.5, and ~1 μM RdRp. The reactions were pre-incubated for 20 min at 30 °C and then started by adding 500 μM UTP, 500 μM ATP, 500 μM CTP, 500 μM GTP. Aliquots of were taken at the time points indicated and stopped with 4 μl formamide loading buffer. Samples subsequently denatured and analysed by 20% denaturing PAGE. The extended capped primers were visualised by phosphorimaging. P-values were determined using an unpaired t-test. To sequence the extended capped primers, products were excised from the 20% denaturing PAGE gel, eluted overnight in water and desalted over NAP-10 columns (GE Healthcare). The isolated RNA was next polyadenylated with polyA polymerase (NEB), reverse transcribed with dTGG primer 5ʹ-CACGACGCTCTTCCGATCTTTTTTTTTTTTTTTTTTGG-3ʹ, and turned into dsDNA using 2^nd^ strand primer 2ND_GA 5ʹ-GTTCAGACGTGTGCTCTTCCGATCTGA+AT+A+CTCAAG-3ʹ (here “+” indicates an LNA base). To remove excess primer, the DNA was treated with exonuclease VII (NEB) for 1 h and subsequently heated to 95 ^o^C for 10 min to denature the exonuclease. Finally, the DNA was amplified with GoTaq (Promega) using primers P5 5ʹ-AATGATACGGCGACCACCGAGATCTACACTCTTTCCCTACACGAC/GCTCTT CCGATCT-3ʹ and i7003 5ʹ-CAAGCAGAAGACGGCATACGAGATACTGGTGTGACTGGAGTTCAGACGTGT/GCTCTTCCGATCT-3ʹ, and TOPO cloned (Invitrogen) for Sanger sequencing. The β-globin mRNA transcription assay was performed as described previously (6).

## Acknowledgements

The authors thank Dr Ervin Fodor for support and suggestions and Dr David Bauer for the i7003 and P5 primers. This work was funded by Wellcome Trust grants 098721/Z/12/Z and 206579/Z/17/Z (to AJWtV), grant 825.11.029 from the Netherlands Organization for Scientific Research (to AJWtV) and an Erasmus+ mobility grant (to JO).

## Conflict of interest

The authors have no competing interests.

## Author contributions

AJWtV designed experiments. JO and AJWtV performed experiments and analysed data. AJWtV wrote manuscript with input from JO.

